# Genetic markers and tree properties predicting wood biorefining potential in aspen (*Populus tremula*) bioenergy feedstock

**DOI:** 10.1101/2021.07.06.450716

**Authors:** Sacha Escamez, Kathryn M. Robinson, Mikko Luomaranta, Madhavi Latha Gandla, Niklas Mähler, Zakiya Yassin, Thomas Grahn, Gerhard Scheepers, Lars-Göran Stener, Stefan Jansson, Leif J. Jönsson, Nathaniel R. Street, Hannele Tuominen

**Affiliations:** Umeå Plant Science Centre (UPSC), Department of Plant Physiology, Umeå University, SE-901 87, Umeå, Sweden; Department of Chemistry, Umeå University, SE-901 87, Umeå, Sweden; RISE AB, Drottning Kristinas väg 61 B, SE-114 28, Stockholm, Sweden; The Forestry Research Institute of Sweden, Ekebo, SE-268 90 Svalöv, Sweden

**Keywords:** Biorefining, feedstock recalcitrance, bioenergy, forest feedstocks, saccharification, biomass

## Abstract

**Background:** Wood represents the majority of the biomass on land and constitutes a renewable source of biofuels and other bioproducts. However, wood is recalcitrant to bioconversion, raising a need for feedstock improvement in production of, for instance, biofuels. We investigated the properties of wood that affect bioconversion, as well as the underlying genetics, to help identify superior tree feedstocks for biorefining.

**Results:** We recorded 65 wood-related and growth traits in a population of 113 natural aspen genotypes from Sweden. These traits included three growth and field performance traits, 20 traits for wood chemical composition, 17 traits for wood anatomy and structure, and 25 wood saccharification traits as indicators of bioconversion potential. Glucose release after saccharification with acidic pretreatment correlated positively with tree stem height and diameter and the carbohydrate content of the wood, and negatively with the content of lignin and the hemicellulose sugar units. Most of these traits displayed extensive natural variation within the aspen population and high broad-sense heritability, supporting their potential in genetic improvement of feedstocks towards improved bioconversion. Finally, a genome wide association study (GWAS) revealed 13 genetic loci for saccharification yield (on a whole tree biomass basis), with six of them intersecting with associations for either height or stem diameter of the trees.

**Conclusions:** The simple growth traits of stem height and diameter were identified as good predictors of wood saccharification yield in aspen trees. GWAS elucidated the underlying genetics, revealing putative genetic markers for bioconversion of bioenergy tree feedstocks.

## Introduction

Lignocellulosic woody biomass represents the majority of biomass on land (Bar-On et al., 2018). This biomass contains mostly three types of natural polymers: cellulose, hemicelluloses and lignin, each of which can be converted into precursors for biofuels and other bioproducts (Percival Zhang, 2013). However, the processes for deconstructing these polymers into usable units remain costly due to structural and chemical hindrance, a problem known as biomass recalcitrance (McCann and Carpita, 2015). Overcoming biomass recalcitrance requires the identification of less recalcitrant feedstocks as well as knowledge on the biological basis of lignocellulose recalcitrance (Van Acker et al., 2014; Wilkerson et al., 2014; Meng et al., 2017; Yoo et al., 2017; Wang et al., 2020). Fast growing trees from the *Populus* genus (poplars, aspens and hybrids) represent promising feedstocks (Mola-Yudego et al., 2017) on account of their lignocellulose composition (Sannigrahi et al., 2010), advanced domestication and efficient cultivation techniques (Dickmann, 2006). Furthermore, genomes of numerous *Populus* species have been sequenced (Tuskan et al., 2006; Ma et al., 2013; Yang et al., 2017; Wang et al., 2018a; Lin et al., 2018; Qiu et al., 2019; Schiffthaler et al., 2019; Hou et al., 2020), which enables investigation of the genetics underlying lignocellulose properties for a better understanding of the biochemistry behind and breeding for less recalcitrance.

Our knowledge of the genetic basis for plant traits has greatly advanced owing to genome wide association studies (GWAS), which relate variation in traits to variation in the sequence of the genomes of different individuals, down to single nucleotide resolution. These variations of nucleotide composition at single loci, also known as single nucleotide polymorphisms (SNPs), can reveal genetic markers for quantitative variation in traits, or even reveal involvement of genes into shaping a quantitative trait (Nordborg and Weigel, 2008).

In a striking example, GWAS of the timing of bud set identified a single locus explaining the majority of local adaptation along a latitudinal gradient in a Swedish population of European aspen *Populus tremula* (Wang et al., 2018a). However, individual loci found by GWAS usually explain only a fraction of the total trait variance, and often a large portion of the genetically heritable variance remains undetermined by significant associations (Nordborg and Weigel, 2008; Du et al., 2018). Nevertheless, finding SNPs associated with only a fraction of the variation in traits of interest could still lead to progress through marker-assisted selection (MAS) or genomics-assisted selection (GAS) for beneficial wood properties (Du et al., 2018).

In *Populus trichocarpa*, GWAS revealed SNPs and genes significantly associated with four wood chemical composition traits (Guerra et al., 2019). Furthermore, associations were discovered between SNPs and 16 wood chemical composition and wood structure traits in *P. trichocarpa* (Porth et al., 2013). Wood chemical composition traits were also linked to SNPs by GWAS in *P. nigra* (Guerra et al., 2013) and *P. deltoides* (Fahrenkrog et al., 2017). Xie et al. (2018) re-evaluated previous associations in *P. trichocarpa* (Porth et al., 2013; Muchero et al., 2015) by focusing on a chromosome known to harbour quantitative trait loci (QTL) for lignin composition, resulting in the identification and characterization of a new transcriptional regulator of lignin biosynthesis. 7 SNPs were identified in association to wood anatomical properties of a *P. trichocarpa* natural population (Chhetri et al., 2020).

Advances in genome (re)sequencing and statistical methods for finding associations in GWAS have facilitated these recent findings (Du et al., 2018; Lin et al., 2018). Yet, the emerging picture of the genetics underlying highly quantitative or complex traits, such as wood properties and bioconversion potential, remains limited, in part due to our limited precision in the quantifications of these traits for entire tree populations (Du et al., 2018; Tuskan et al., 2019). For example, lignin is composed of different types of monomers, and measurement of only the total amount of lignin in wood obscures the influence and the regulation of the abundance of the different types of lignin monomers (Tuskan et al., 2019). Increasing the number of analysed traits and the depth of the analyses is likely needed for GWAS analyses of especially the complex traits (Ingvarsson and Street 2011; Du et al., 2018; Tuskan et al., 2019). Extensive phenotyping also allows better characterization of the relationships between traits, for example to identify which wood chemical composition and structure traits determine wood bioconversion potential.

Here, we present a large-scale phenotyping effort, monitoring 65 traits related to wood properties, tree growth, and wood saccharification in a common garden trial comprising a collection of natural aspen (*Populus tremula*) genotypes (the so-called SwAsp collection) collected across Sweden (Luquez et al., 2008). Through genetic correlation and multivariate analyses, we identified wood chemical composition and structural traits correlating with recalcitrance as well as whole stem bioconversion potential. Through GWAS, we identified several novel genetic loci linked to both tree growth and whole stem bioconversion potential.

## Results

### Natural variation in 65 growth, wood, and biorefinery traits in aspen

Natural variation in wood and biorefinery traits was investigated in 113 clonally replicated aspen trees of the SwAsp collection. After ten years of growth in a common garden in southern Sweden, we measured stem height and diameter, wood chemical composition (20 traits), wood structural and anatomical properties (17 traits), as well as recovery of monosaccharides from wood saccharification with or without acidic pretreatment (25 traits), amounting to 64 traits (Fig. 1a, Additional file 1). Finally, we estimated total wood glucose yield (TWG; Additional file 1). While glucose release provides information about biomass recalcitrance to saccharification, TWG provides a proxy for overall tree performance.

**Figure 1.**
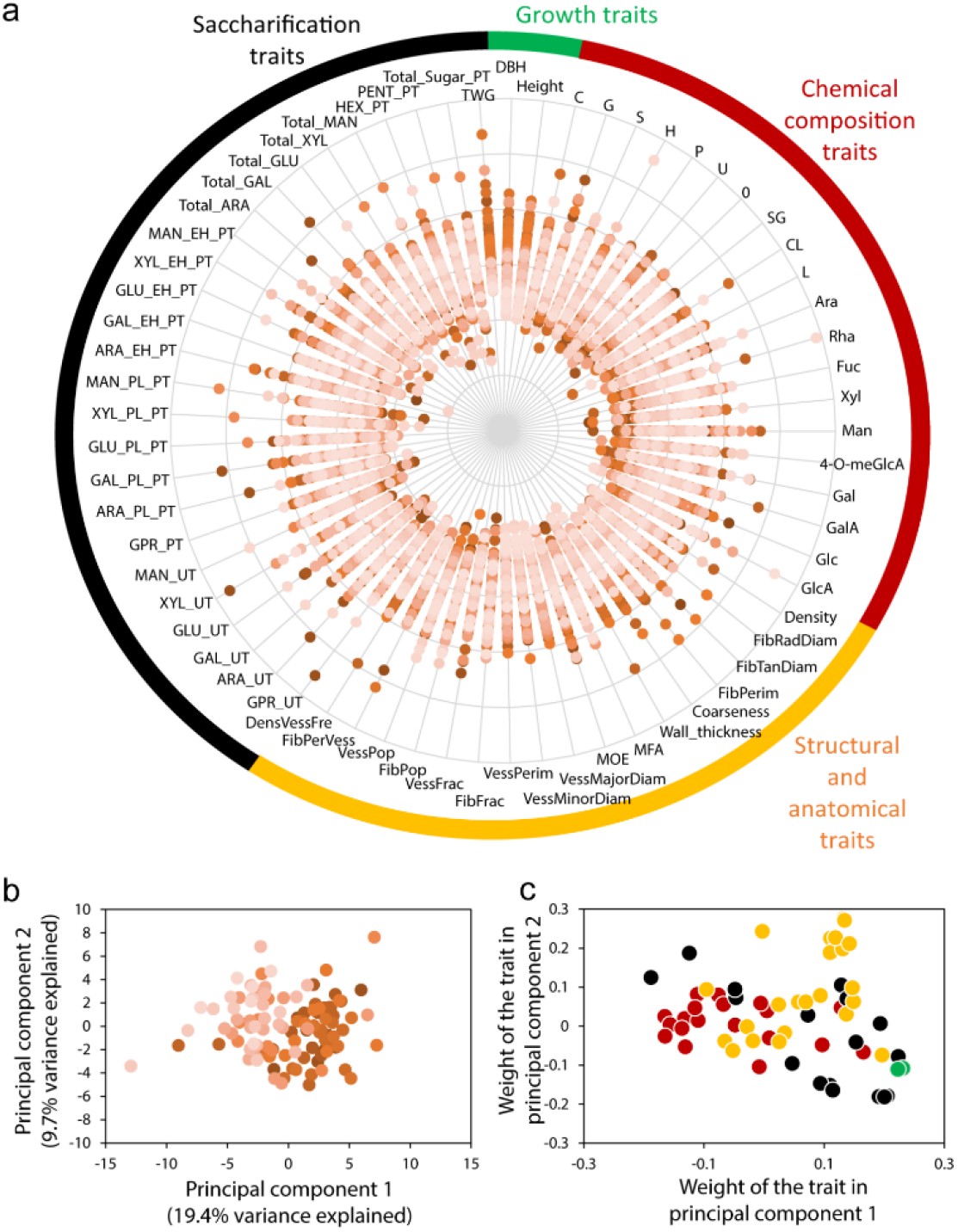
The SwAsp natural variants display a wide range of phenotypic variation. **(a)** 65 traits related to tree growth, wood chemical composition, wood structure and anatomy, and saccharification traits of woody biomass in 10-year-old aspen trees from 113 genotypes. Each point represents the median scaled and centered measurement of a trait for one genotype (z-transformation across the tree population for each trait). Colored labels around the plot indicate categories of traits (chemical composition; structure and anatomy; saccharification and growth). Abbreviations are defined in Additional file 1. **(b)** Principal Component Analysis (PCA) scatter plot showing that the SwAsp genotypes differ from each other based on their wood properties and saccharification. Colors indicate the 12 different locations of origin for the different genotypes in Sweden. **(c)** Coefficients scatter plot of traits. Each point corresponds to a trait; while the colors indicate which trait category they belong to (as in (**a**)).

All traits showed phenotypic variation among the genotypes (Fig. 1a, Additional file 1). Around 30 % of the total variation was explained by the two first components in a principle component analysis (Fig. 1b), with these largely being influenced by variation in the saccharification traits (Fig. 1c). Indeed, the saccharification traits, such as glucose release after enzymatic hydrolysis with pretreatment and total wood glucose yield, displayed almost 50% increase from the lowest to the highest yielding genotype (Fig. 2, Additional file 1). Lignin traits, such as total lignin content and the ratio between the syringyl (S) and guaiacyl (G) type lignin (SG), that are central in determining feedstock recalcitrance, also varied substantially among the different genotypes (Fig. 2, Additional file 1).

**Figure 2.**
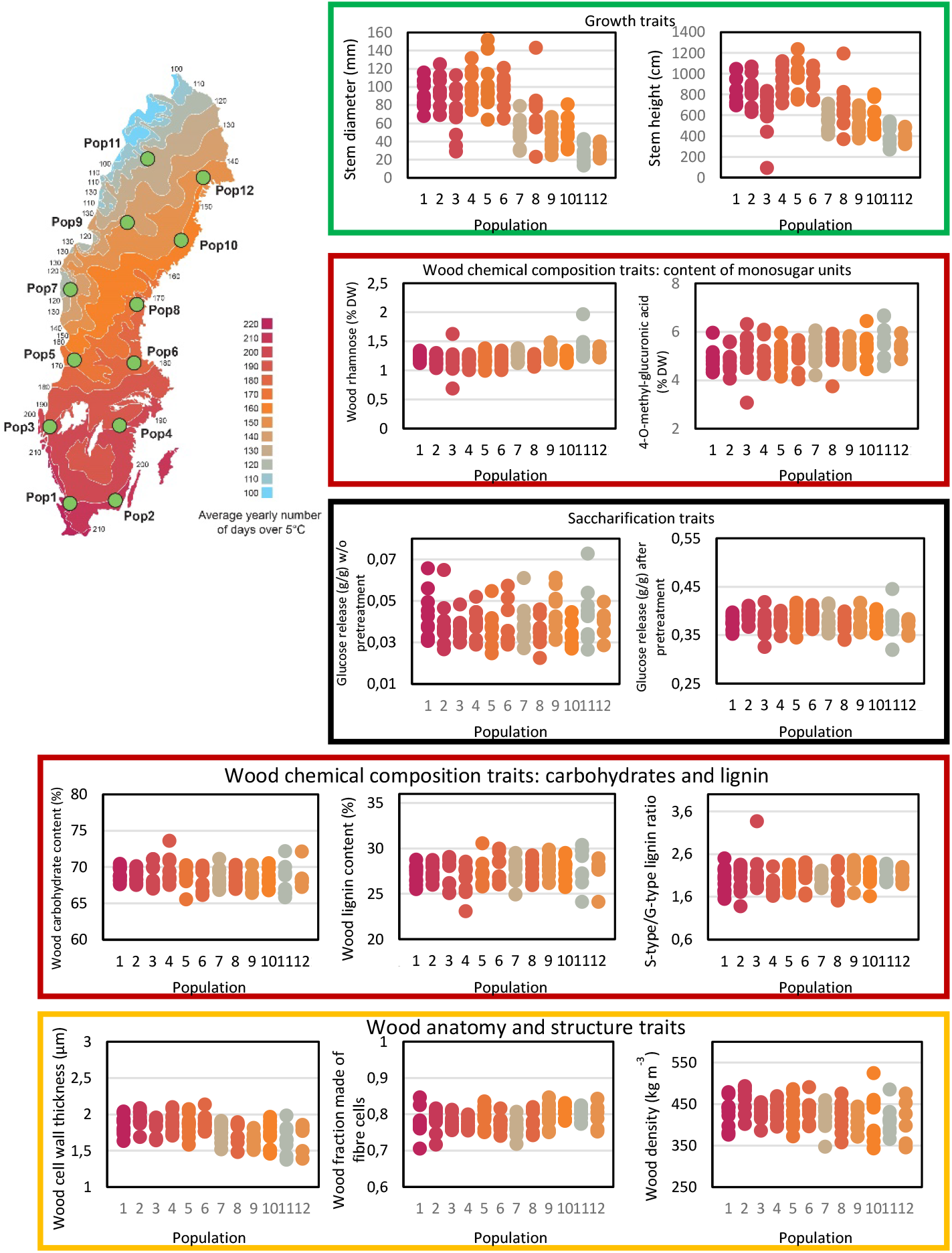
Wood and biorefining traits and geographical origin of the SwAsp trees. The average values are shown for key representative traits for growth, wood chemical composition, wood anatomy and structure, and traits related to saccharification for the 113 different genotypes of the SwAsp collection. The values are grouped according to the geographic origin of the genotypes in 12 locations across Sweden. The locations for the different geographic origins (Pop1-Pop12) are illustrated on the map in the upper left corner.

We estimated the broad sense heritability (H^2^) of the different traits (Additional file 2). Some traits, such as those linked to wood xylose units and xylose released by saccharification, showed nearly no heritability, while traits related to tree growth and wood anatomy showed moderate to high heritability (H^2^ > 0.5). Wood chemical composition traits showed varying heritability; generally lower for wood monosaccharide units and higher for lignin composition traits, especially the S-type and G-type lignin content (Additional file 2).

Next, genetic correlations were estimated among the different traits. The tree growth traits (height and diameter) correlated positively with wood density, xylem cell wall thickness, xylem cell diameters and wood carbohydrate content (Fig. 3, Additional file 3). The correlations for the saccharification traits varied somewhat depending on the sugar analysed, but the release of sugars having the highest abundance in wood, glucose (GLUEHPT) and xylose (XYLEHPT), correlated positively with the growth traits of the trees and negatively with lignin content. Another striking result was that the glucose release GLUEHPT correlated negatively with the wood content of all hemicellulose sugar units (Ara, Fuc, Gal, GalA, GlcA, Man, 4-O-meGlcA, Rha, Xyl) (Fig. 3, Additional file 3). A slight positive correlation was present between GLUEHPT and the S to G lignin ratio (SG).

**Figure 3.**
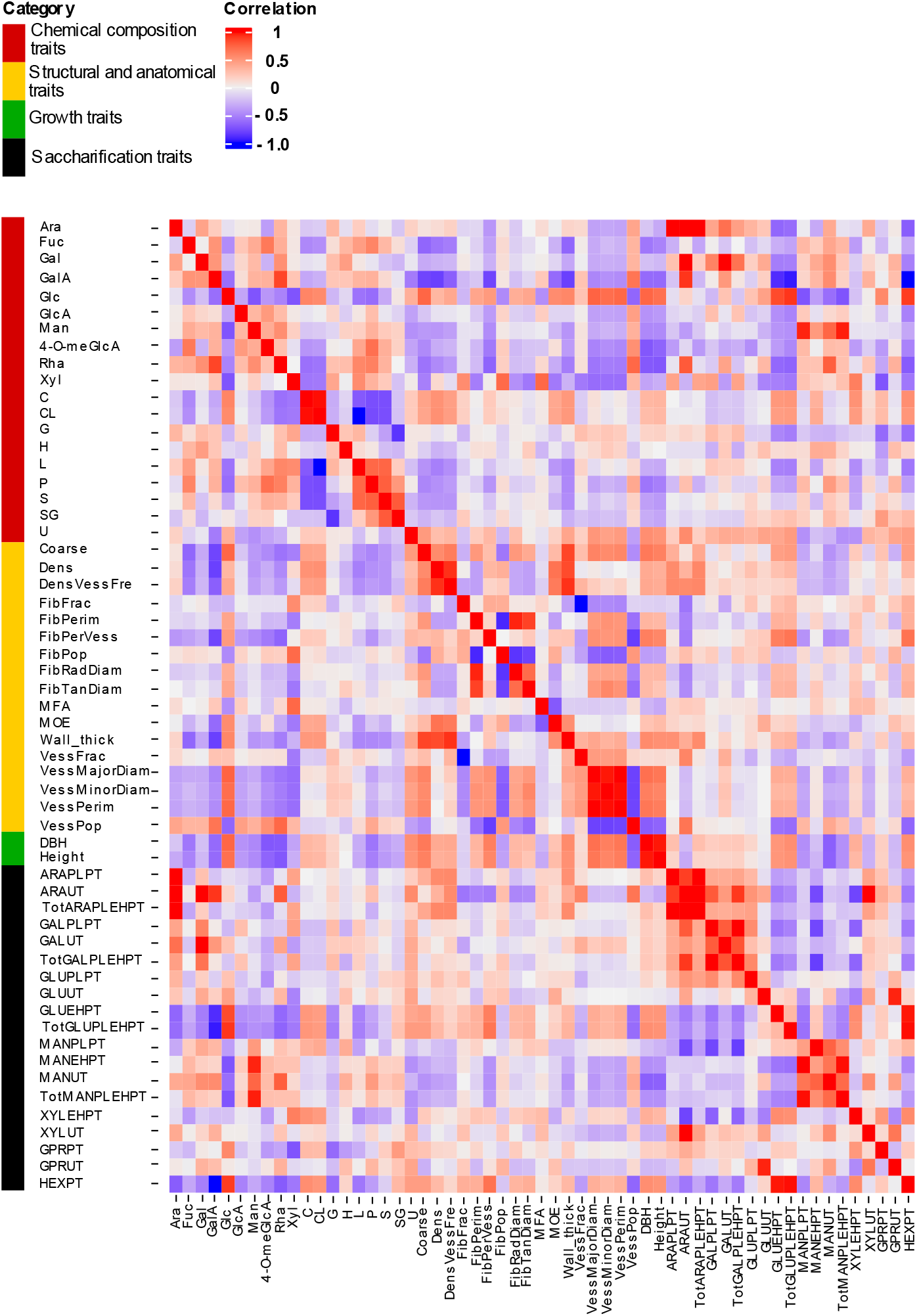
Pairwise genetic correlations between 58 traits in SwAsp trees. The vertical sidebar represents the four categories of traits: wood chemical composition (red), wood structure and anatomy (yellow), growth (green) and wood saccharification (black). Six analysed SwAsp traits were omitted from the correlation analyses due to very low heritabilities. Trait abbreviations are defined in Additional file 1.

In a phenological study of the SwAsp population, the timing of bud set was shown to correlate with the geographical origin of the genotypes (Wang et al., 2018a). On the other hand, studies of secondary metabolites or leaf shape in that same population showed no correlation between these traits and the geographical origin of the genotypes (Keefover-Ring et al., 2014; Mähler et al., 2020). These previous observations raise the question of whether wood and biorefinery traits display a geographical cline. The growth traits, stem height and diameter, showed an expected, clear relationship to the geographical origin of the genotypes (Fig. 2, Additional file 1). Even though the traits related to wood chemical composition, wood anatomy and structure, and saccharification did not show clear geographic clines on a population level (Fig. 2), correlation analysis on the clonal basis showed significant effect of the geographic origin for several traits. For instance, relative carbohydrate content, cell wall thickness, vessel diameter and wood density correlated negatively with the latitude of the clonal origin, while relative lignin content, content of hemicellulose sugar units and S/G lignin ratio correlated positively with the latitude (Additional file 1). Since the heritabilities for these traits were high (Additional file 2), these results suggest that aspen genotypes from northern Sweden had, on average, more total lignin, S-type lignin and hemicelluloses, and less carbohydrates, lower vessel diameter and wood density than genotypes from southern Sweden.

### Identification of traits that influence wood recalcitrance

To better characterize the traits influencing wood recalcitrance to bioprocessing, we performed multivariate analyses for the glucose release from saccharification, as well as for the TWG. We employed orthogonal projections to latent structures (OPLS; Trygg and Wold, 2002), which considers all traits simultaneously, to get an overview of the relationships between wood properties and glucose release or TWG (Fig. 4). OPLS models were created that explained high proportion of the variation for both the glucose release after enzymatic hydrolysis with pretreatment (GLUEHPT) and for TWG (R^2^ = 0.56 and 0.52, respectively), but the predictivity of the model was not strong (Q^2^ = 0.17 and 0.29, respectively). The OPLS models supported negative contribution of wood hemicellulose sugar units and lignin on both GLUEHPT and TWG, while several wood anatomy traits, such as diameter of the fibers and the vessels, the ratio of fibers to vessels (FibPerVess) and coarseness (weight of fibers over a certain length of wood), contributed positively to the models of both traits (Fig. 4).

**Figure 4.**
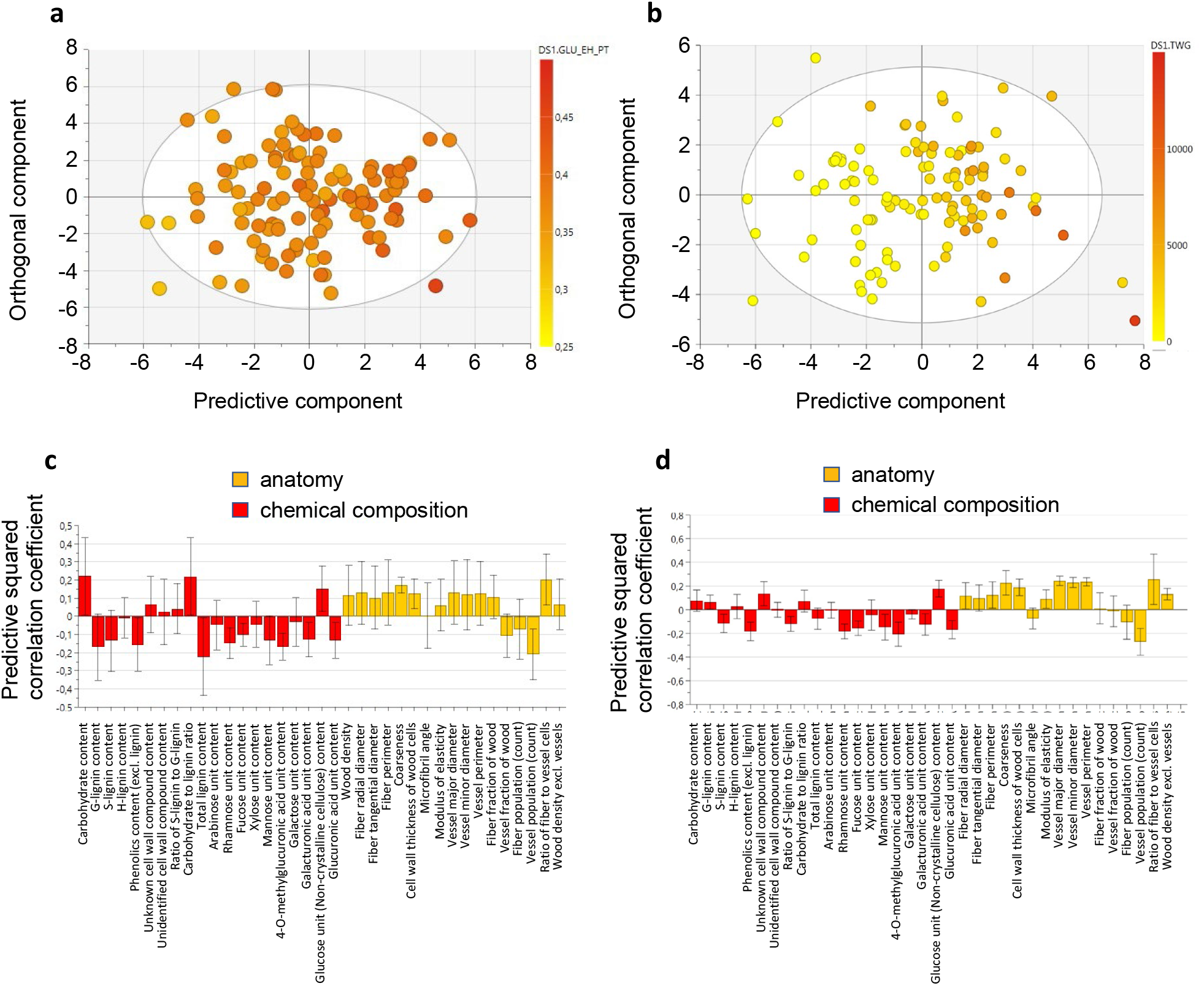
Multivariate analysis of the relationships between wood properties and glucose release or TWG. **(a**,**b)** Orthogonal Projection to Latent Structure (OPLS) scatter plot showing separation in glucose release after pretreatment **(a)** and total wood glucose yield **(b)**. The points on the scatter plot correspond to SwAsp genotypes; while their color indicates the median for the trait in each genotype. **(c**,**d)** OPLS loadings plot for glucose release after pretreatment **(c)** and total wood glucose yield **(d)** in relation to wood chemical composition and wood anatomy traits. The bars indicate the coefficient (“weight”) of each trait in the OPLS model. The traits with positive values correlate positively and the traits with negative values negatively with glucose release after pretreatment **(c)** and total wood glucose yield **(d)**.

### Genetic polymorphisms are significantly associated with total wood glucose yield

To further decipher the genetics underlying wood properties and amenability to improvements in bioprocessing, we performed a genome wide association study (GWAS). In this analysis, latitude of origin of each SwAsp genotype was included as a covariate due to the presence of the latitudinal clines (Additional file 1). An FDR cutoff of 0.1 was selected to identify putative associations (Storey and Tibshirani, 2003).

Single nucleotide polymorphisms (SNPs) with FDR value less than 0.1 (*q*-value < 0.1) were identified in 17 loci for five traits (Table 1, Additional file 4). Most of the SNPs were located in intergenic regions or upstream/downstream of gene coding regions.

**Table 1.** Genes and genomic features associated with SNPs at *q*-value < 0.1 in the SwAsp genome-wide association study of 65 traits monitored in the Swedish aspen collection.

| Gene <sup>1</sup> | Feature <sup>2</sup> | Description(s) | The number of SNPs associated with each trait |  |  |  |  |  |
| --- | --- | --- | --- | --- | --- | --- | --- | --- |
|  |  |  | DBH | Wall thickness | FibFrac | Height | VessFrac | TWG |
| 7c16159;<br>7c16161 | Intergenic |  | 1 |  |  |  |  | 1 |
| 15c29048;<br>15c29049 | Intergenic |  | 1 |  |  | 1 |  | 1 |
| 9c19049;<br>9c19048 | upstream;<br>downstream | Oxa2A membrane insertase;<br>hypothetical protein | 1 |  |  |  |  | 1 |
| 3c8002 | Downstream | MATE efflux family protein | 1 |  |  |  |  |  |
| 1c2354;<br>1c2355 | Intergenic |  |  |  |  | 18 |  | 12 |
| 4c9179;<br>4c9180 | Intergenic |  |  |  |  | 2 |  | 2 |
| 5c11372;<br>5c11373 | Intergenic |  |  |  |  | 1 |  | 1 |
| 10c20558 | Exonic | 2-oxoglutarate dehydrogenase, E1 |  |  |  |  |  | 1 |
| 10c21196;<br>10c21197 | upstream;<br>downstream | Geranyl diphosphate synthase 1;<br>hypothetical protein |  |  |  |  |  | 1 |
| 16c29591 | UTR3 | L-galactono-1,4-lactone dehydrogenase |  |  |  |  |  | 1 |
| 17c31917;<br>17c31919 | Intergenic |  |  |  |  |  |  | 1 |
| 18c33227;<br>18c33228 | Intergenic |  |  |  |  |  |  | 1 |
| 10c22263;<br>10c22264 | Intergenic |  |  |  |  |  |  | 1 |
| 14c26895 | Intronic | AGAMOUS-like 20 |  |  |  |  |  | 1 |
| 3c6553;<br>3c6554 | Intergenic |  |  | 6 |  |  |  |  |
| 12c24540;<br>12c24542 | upstream; UTR3-downstream | DnaJ homolog subfamily B member;<br>Major facilitator superfamily protein |  |  | 5 |  | 5 |  |
| 17c31483 | Downstream | NA |  |  |  |  | 1 |  |

<sup>1</sup> The full name of the *P. tremula* gene models includes “Potra2n” in front of the “gene”.
<sup>2</sup> The feature upstream and downstream indicates location of the SNPs within 2 kbp from the coding region, while the feature intergenic indicates on SNPs further than 2 kbp from the coding region.

No significant associations were observed for glucose release rates, but 11 associations for TWG with *q*-value < 0.1 were identified on 11 chromosomes, with each individual SNP explaining 22 to 26 % of phenotypic variation (Table 1, Additional file 4). Six of the loci for TWG associations intercepted with loci containing SNPs for either stem diameter at breast height (DBH) or stem height (Height) (Table 1, Additional file 4). These six loci were all intergenic except for chr9_2882991_T_C which was located 362 bp upstream from Potra2n9c19049 (Oxa2A membrane insertase) and 757 bp downstream from Potra2n9c19048 (hypothetical protein) (Table 1, Additional file 4). The chr9_2882991_T_C SNP also showed statistically significant differences in the phenotypes between the SwAsp genotype groups with homozygous and heterozygous alleles for not only TWG but also DBH and height (Fig. 5b-d). While the proportion of phenotypic variation explained by chr9_2882991_T_C was 0.26, the minor allele frequency was low (0.05) such that there was only one SwAsp individual with homozygous minor alleles for this SNP and the TWG phenotype was absent for this individual (Fig. 5d). In addition to the six loci intercepting with the DBH and/or height, GWAS revealed a locus with 12 TWG-SNP associations with *q*-value < 0.1 (Fig. 5a) in an intergenic region in the chromosome 1, in a region spanning 3327 base-pairs and including 38 SNPs with R^2^ values > 0.2, considered to be in linkage disequilibrium with the 12 significant SNPs (Additional file 4). The most significant SNP by *P-*value and *q*-value in this locus was chr1_28056992_G_A (with a major allele frequency of 0.132 and PVE of 0.26). The SwAsp genotype groups with homozygous and heterozygous alleles for the chr1_28056992_G_A significantly partitioned the variance of TWG as well as DBH and height (Fig. 5e-g). Out of the remaining putative associations for TWG, the only SNP that resided in the coding region of a gene, chr10_2830421_T_G, corresponded to Potra2n10c20558 (E1 subunit of oxoglutarate dehydrogenase). Although this result is based on only one SwAsp genotype with the homozygous recessive (GG) allele (Fig. 5h), it also showed statistically significant differences in the height and DBH phenotypes among the allele groups (Fig. 5i and j).

**Figure 5.**
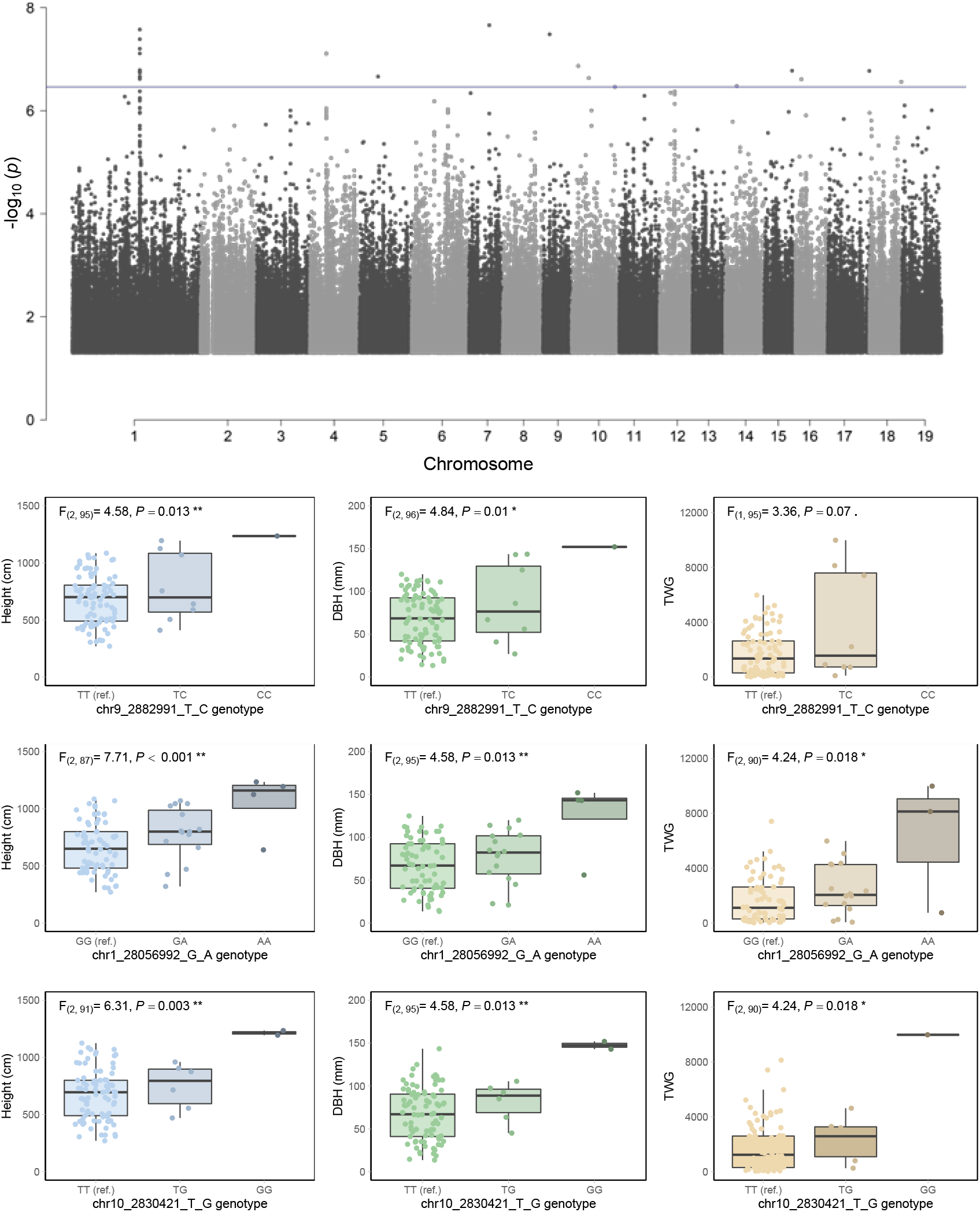
Genome wide association analysis of total wood glucose yield (TWG). **(a)** Manhattan plot for total wood glucose yield (TWG). Each point indicates location of a SNP along the 19 chromosomes of *Populus tremula*. The blue horizontal line indicates the *q*-value level of 0.1. The least significant SNPs (*P*-values > 0.05) have been omitted for plot clarity. **(b-j)** Tree height, diameter at breast height (DBH) and TWG in relation to their SNP genotype for the three most significant associations for TWG. Boxplots show phenotypic values of height, diameter and TWG amongst allele classes for the SNPs with the smallest p-values in the association tests; chr1_28056992_G_A had the highest statistical significance among the SNPs in the chromosome 1 GWAS hotspot for TWG (see also Additional file 4). The jittered points around each box represent median phenotypic values of the SwAsp clonal replicates. Analysis of variance F-ratios and *P*-values are reported, where the dependent variable is the phenotype and the and independent variable is the SNP genotype class, and significance indicated at. <0.1, * <0.05, and ** < 0.01. The three traits are shown for the SNPs chr9_2882991_T_C **(b-d)**, chr1_28056992_G_A **(e-g)** and chr10_2830421_T_G **(h-j)**.

SNPs with *q*-value <0.1 were found for the fraction of the wood made of fibers (FibFrac) and vessels (VessFrac) downstream of Potra2n12c24540 (DnaJ homolog subfamily member) and upstream of Potra2n12c24542 (Major facilitator superfamily protein member) (Table 1, Additional file 4). Furthermore, six SNPs with *q*-value <0.1 were identified for cell wall thickness of the wood cells (Wall thickness) in an intergenic region.

## Discussion

Wood biomass from fast growing trees represents a promising source of biofuels and other bioproducts in the forthcoming transition away from fossil fuels (Percival Zhang, 2013; Ragauskas et al., 2014). The high cost of deconstructing woody biomass, however, hinders wood biorefining (McCann and Carpita, 2015). To overcome this biomass recalcitrance, it is necessary to understand how wood properties relate to wood recalcitrance. We report here the phenotyping of a population of aspen genotypes for 65 traits related to tree growth, wood anatomy and structure, wood cell wall chemical composition, and wood bioprocessing yield.

Using genetic correlations and multivariate modelling, we identified a set of wood traits that correlate with the glucose yield from saccharification. Lignin content and especially G-lignin content had a negative influence on the glucose yield after enzymatic hydrolysis with pretreatment, which is in line with a positive effect of S/G ratio in our earlier analysis of 40 transgenic *Populus* lines (Escamez et al., 2017), as well as in the analyses of *P. trichocarpa* (Studer et al., 2011; Yoo et al., 2017) and *Salix viminalis* (Ohlsson et al., 2019) natural variants. A negative effect of S/G ratio was reported in a small selection of natural *P. trichocarpa* variants, and it was proposed that S/G ratio might instead influence xylose release after enzymatic hydrolysis (Meng et al., 2017). We could not confirm this as no correlation was found between xylose release after enzymatic hydrolysis and S/G ratio in our dataset (Fig. 3). In addition to lignin, a consistent negative influence on sugar (glucose) yields after saccharification with pretreatment was imposed by the hemicellulose sugars (Fig. 3 and 4). This is most probably related to the fact that the pretreatment was adjusted to a rather mild level of severity, resulting in part of the hemicelluloses remaining intact in the feedstock: the wood xylose unit content was 0.2-0.3 g/g DW depending on the clone, while 0.12-0.16 g/g DW xylose was retrieved from the biomass into the pretreatment liquid (Additional file 1). Hemicelluloses are, in addition to lignin, the most important wood recalcitrance factors (Martín et al., 2022). It is therefore likely that the hemicelluloses retained in the wood after the pretreatment limited the saccharification efficiency. Furthermore, the contents of the hemicellulose sugars also correlated in a similar, negative fashion with the relative carbohydrate content of wood (Fig. 2), which could also contribute to the negative influence of the hemicellulose sugars on the glucose release.

Identifying the genetics underlying wood properties that foster bioprocessing potential help with selecting or creating superior biorefinery feedstocks (Fahrenkrog et al., 2017). GWAS has frequently been used to identify single nucleotide polymorphisms associated with wood properties (Porth et al., 2013; Tuskan et al., 2019; Chhetri et al., 2020). GWAS for saccharification traits is rare in forest tree species, but a notable association was found in *Salix viminalis* for glucose release in a non-coding region of the genome (Ohlsson et al., 2019). We found no associations for glucose release, but several loci of putative associations with the total wood glucose yield (Table 1). One of these was located in the coding region of E1 subunit of oxoglutarate dehydrogenase (Potra2n10c20558) which participates in the mitochondrial citric acid cycle to provide reducing power for oxidative phosphorylation and carbon skeletons for various metabolic pathways. The importance of this enzyme is supported by a dramatic influence of the two homologous *Arabidopsis* genes (AT3G55410 and AT4G26910) as well as the E2 subunit of the same complex on growth and biomass production in *Arabidopsis* (Condori-Apfata et al., 2019; 2021). Interestingly, the Oxa2A membrane insertase (Potra2n9c19049) and the geranyl diphosphate synthase (Potra2n10c21196), located in close promixity of a SNP for TWG, are both mitochondrial proteins (Ducluzeau et al., 2012; Kolli et al., 2019), supporting the link between mitochondrial function and TWG yield.

TWG is a composite trait consisting of glucose yields after saccharification on a whole-tree-biomass basis. Consequently, we identified six polymorphic loci that intercepted for TWG and the biomass-related parameters of tree height and stem diameter (Table 1), pointing out the importance of tree biomass yields on TWG. For breeding purposes, it is an interesting question which one is more important for saccharification yields on a whole tree basis; biomass production or saccharification yields (per g material). In an earlier study of transgenic hybrid aspen trees, we found that increased biomass production compensated for the loss of glucose release (per gram) caused by the transgene expression (Gandla et al., 2021). Vice versa, gains from increased glucose release after saccharification of lignocellulosic feedstocks have frequently been offset by decreased biomass production of trees (Bonawitz and Chapple, 2013; Vanholme et al., 2013; Van Acker et al., 2014; Ha et al., 2021). It therefore seems that saccharification yields can be efficiently increased simply by increasing biomass production of the trees. Biomass production is influenced by tree volume and wood density, and it was interesting that in the currently investigated population of aspen trees it was the tree volume-related traits of height and diameter that correlated better with the glucose release after saccharification with pretreatment than wood density (Fig. 3), and that tree height and diameter were the traits that intercepted with TWG in the GWAS analysis (Fig. 5). This implies that the simple measurements of tree height and diameter might be sufficient to predict saccharification yields on a whole-tree basis. Earlier studies have seldom approached this question since saccharification has traditionally been defined on a process basis, resulting in identification of chemical composition as the most important factor influencing sugar yields. However, a very similar conclusion was drawn in a recent analysis of field grown *P. trichocarpa* trees where stem diameter was identified as the main driver for ethanol yield on a whole-field basis (Happs et al., 2021). This leads to the question of what is the impact of tree volume on the other traits determining biomass production or saccharification? Notably, positive genetic correlation existed in our population between the tree volume traits and wood density as well as glucose release, which both act to increase the TWG (Fig. 3). Furthermore, negative correlation existed between tree growth and lignin content and hemicellulose sugars, implying that breeding efforts towards increased tree volume production might, similar to the currently investigated material, supress the accumulation of wood chemical properties that have negative influence on both glucose release rate and TWG.

## Conclusions

We identified significant natural variation in growth and wood-related traits in aspen, which allowed identification of chemical and genetic markers for bioprocessing purposes of lignocellulosic feedstocks. Our data indicate that whole tree saccharification yields can be improved, at least in *Populus* feedstocks, by simply breeding for increased tree volume growth without a negative impact on wood parameters, such as wood density or the content of lignin and hemicelluloses, that also influence saccharification efficiency and yield.

## Materials and methods

### Plant material

The Swedish Aspen (SwAsp) collection consists of 113 *Populus tremula* aspen genotypes from 12 locations across Sweden (Luquez et al., 2008). The genotypes represent potential sub-populations (Fig. 2), but whole-genome sequencing and sequence comparisons have shown that these genotypes are mostly unrelated (Wang et al., 2018a).

The genotypes were clonally propagated in 2003 from root cuttings and grown in a randomized block experiment in a plantation in southern Sweden (Ekebo, 55.9°N). Three to five trees per genotype were successfully established in 2004 (Luquez et al., 2008; Wang et al., 2018a).

After ten years of growth, tree height and diameter at breast height (DBH) were measured, and wood samples were collected from the stem. At 79 cm above ground, a 1-cm-thick section of the stem was collected, and the south-western facing quarter of the stem section was aliquoted for wood chemical composition analyses. In addition, 80-90 cm above ground, another piece of stem was harvested for analysis of wood anatomical and structural properties from the south-western facing quarter of the stem section. We obtained a full set of successful phenotypic measurements for a total of 418 trees, representing two to five replicates per genotype (Additional file 1).

### Analyses of wood chemical composition

The wood quarters selected for compositional analyses were manually debarked, cut into roughly match-stick-sized wood pieces and freeze dried (CoolSafe Pro 110-4, LaboGene A/S, Denmark). This material was homogenized by coarse milling (Retsch ZM 200 centrifugal mill, Retsch GmbH, Germany) and sieved (Retsch AS 200) into two particle size fractions. The fraction of particle size between 0.1 mm and 0.5 mm was aliquoted for subsequent saccharification experiments (see below), while the fraction of particle size under 0.1 mm was aliquoted for pyrolysis coupled with gas chromatography followed by mass spectrometry analysis (pyrolysis-GC/MS) and monosaccharide composition analysis. Both analyses were performed as technical duplicates for each tree.

Carbohydrate content, lignin content, lignin composition, and content of other phenolics were determined by pyrolysis-GC/MS as previously described (Gerber et al., 2016). Briefly, 40 μg - 80 μg of homogenized wood powder was loaded into an autosampler (PY-2020iD and AS-1020E, Frontier Labs, Japan), allowing a sub-sample (∼1 μg) into the pyrolizer of the GC/MS apparatus (Agilent, 7890A/5975C, Agilent Technologies AB, Sweden). Following pyrolysis, the samples were separated along a DB-5MS capillary column (30 m × 0.25 mm i.d., 0.25-μm-film thickness; J&W, Agilent Technologies), and scanned by the mass spectrometer along the m/z range 35 – 250. The GC/MS data were processed as previously described (Gerber et al., 2012). Results were normalized by expressing the area of each peak as a percentage of the total peak area considering all peaks.

Cell wall monosaccharide units were quantified following the acidic methanolysis and trimethylsilyl (TMS) derivatization method as described previously (Gandla et al., 2015). Briefly, wood powder was washed with HEPES buffer (4 mM, pH 7.5) containing 80% ethanol, as well as methanol:chloroform 1:1 (V:V) and acetone to generate alcohol insoluble residues (AIRs) which were then dried. To avoid contamination with glucose from starch, the AIRs were treated with 1 unit per AIR mg of type I α-amylase (Roche 10102814001, Roche GmbH, Germany). The de-starched AIRs, and inositol as an internal standard, were methanolysed using 2 M HCl/MeOH at 85°C for 24 h. Following repeated washes with methanol, the samples and standard were silylated using Tri-sil reagent (3-3039, SUPELCO, Sigma-Aldrich, Germany) at 80°C for 20 min. The solvent was evaporated under a stream of nitrogen and pellets were dissolved in 1 mL hexane and filtered through glass wool. The filtrates were evaporated until 200 μL remained, of which 0.5 μL were analysed by GC/MS (7890A/5975C; Agilent Technologies AB, Sweden) according to Sweeley et al. (1966). The levels of the sugars and sugar acids are presented in the hydrous form.

### Saccharification assays and total wood glucose yield (TWG)

Saccharification assays without or with acid pretreatment of the biomass were performed following an established methodology (Gandla et al., 2015). In short, 50 mg of dry wood powder (moisture measured with an HG63 moisture analyser, Mettler-Toledo, USA) with particle size between 0.1 mm and 0.5 mm were pretreated with 1% (w/w) sulphuric acid (fraction of sulphuric acid based on the mass of the whole reaction mixture) during 10 min at 165°C in a single-mode microwave system (Initiator Exp, Biotage, Sweden), or remained untreated. The pretreated samples were centrifuged to separate the solid fraction from the pretreatment liquid. The solid fraction was washed with ultrapure water and sodium citrate buffer (50 mM, pH 5.2). The washed, pretreated solid fraction as well as the untreated samples were enzymatically hydrolysed 72 h at 45°C under agitation, using 25 mg of a 1:1 (w/w) mixture of liquid enzyme preparations Celluclast 1.5 L (measured CMCase activity of 480 units per gram of liquid enzyme preparation, following Ghose (1987)) and Novozyme 188 (measured β-glucosidase activity of 15 units per gram liquid enzyme preparation, following Mielenz (2009) (Sigma-Aldrich). Sodium citrate buffer (50 mM, pH 5.5) was added to reach 1 g of final reaction mixture. During enzymatic saccharification, samples were collected at 2 h and 72 h. Glucose production rates were determined at 2 h using an Accu-Chek ®Aviva glucometer (Roche Diagnostics Scandinavia AB, Sweden). Monosaccharide (arabinose, galactose, glucose, xylose and mannose) yields in pretreatment liquids and enzymatic hydrolysates collected at 72 h were determined using high-performance anion-exchange chromatography with pulsed amperometric detection (Ion Chromatography System ICS-5000, Dionex, USA) as previously described (Wang et al., 2018b). Saccharification was performed on technical duplicates for each tree.

The total-wood glucose yield from an entire tree trunk (TWG) was calculated using the formula TWG = 1/3 × π × height × (diameter/2)^2^ × wood density × glucose release_(AFTER PRETREATMENT)_, as previously described (Escamez et al., 2017), assuming a conical shape of the tree stem.

### Anatomical and structural characterisation

Anatomical and structural features were determined on parallelepipedal wood pieces across the stem diameter using the SilviScan® instrument (CSIRO, Australia) which consists of three separate units: (i) a cell scanner with a video microscope for measurement of the numbers and sizes of fibres and vessels, (ii) a density scanner recording X-ray absorption images for measuring wood density, and (iii) a diffraction scanner recording X-ray diffraction images for measuring the microfibril angle. The measurements on these parallelepipedal wood pieces were then projected onto the entire wood section to reflect the average values for the entire wood section of each tree. Full description of the different traits from the SilviScan measurements can be found in Additional file 1.

### Statistical estimations of the genetic parameters

The genetic parameters for each trait were estimated statistically based on measurements on individual trees for each genotype according to the model Y_ijk_ = μ + b_i_ + c_j_ + e_ijk_ where Y_ijk_ is the observation k in block i for clone j, μ is the mean of the trait in this trial, b_i_ is the fixed effect of block i, c_j_ is the random effect of clone j (normally and independently distributed with mean 0 and variance V_c_; NID[0,V_c_]), and e_ijk_ is the random error term for observation ijk (NID[0,V_e_]). The variances V_c_ and V_e_ were estimated for each trait according to the Restricted Maximum Likelihood (REML) method using the ASREML software (Gilmour et al., 1997). To estimate genetic parameters, we considered that V_c_ is equal to V_G_ (the genotypic variance among clones for the trait) and V_e_ is equal to V_E_ (the environmental variance for the trait). Correlation analysis was not performed for traits ARA_EH_PT, GAL_EH_PT, Total_SUGAR_PT, Total_XYL_PL+EH_PT, XYL_PL_PT and PENT_PT due to very low heritabilities.

For each trait, broad-sense heritability (*H*^*2*^) was estimated by dividing genotypic variance (V_G_) by the total variance of this trait V_T_ where V_T_ = V_G_ + V_E_. The genotypic coefficient of variation (CV_G_) for a trait was calculated by dividing the genotypic standard deviation of the trait 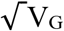 by the mean value of the trait 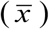, and multiplying the result by 100. The genetic correlation (r_G_) between trait 1 with genotypic variance V_G1_ and trait 2 with genotypic variance V_G2_ was calculated by dividing the genotypic genetic covariance (cov_G1G2_) between these traits by the square root of the product of their individual genetic variances; 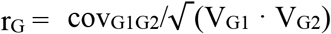.

### Multivariate analyses

Multivariate analyses using all wood traits to predict glucose release by saccharification, or total wood glucose yield (TWG), were performed using Orthogonal Projections to Latent Structures (OPLS) regression (Trygg and Wold, 2002), with 1 + 3 components.

### Genome-wide Association Study (GWAS)

Phenotypic data were subjected to a scripted pipeline, comprising a set of quality control steps and the estimation of a best linear unbiased predictor (BLUP) phenotypic value for each SwAsp genotype and for each trait in the GWAS. The pipeline is described in Schiffthaler et al. (2019) and scripts are available at https://github.com/sarawestman/Genome_paper. Briefly, phenotypic outliers were removed using the ‘OutlierTest’ function of the ‘car’ package in R (R Core Team, 2022; version 3.0.10; Fox & Weisberg, 2019), phenotypes were tested with the Shapiro-Wilk test and any non-normally distributed random effects or error terms were transformed using an Ordered Quantile normalization in the ‘bestNormalize’ package in R (version 1.6.1, Peterson & Cavanaugh, 2019). Subsequently, a BLUP with a restricted maximum likelihood approach was used to estimate the genotypic effect of a given phenotype, as detailed in Wang et al. (2018a) using the model:

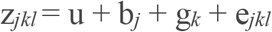

Where z_jkl_ is the phenotype of the *lth* individual in the *jth* block from the *kth* genotype, u is the grand mean and e_jkl_ is the residual error term. The genotype and residual terms were considered random effects, and field block was considered a fixed effect.

Details of the SwAsp DNA sequencing and SNP calling have been described previously (Rendón-Anaya et al., 2021), resulting in 99 unrelated individual genotype sequences for GWAS with biallelic, high quality sites (single nucleotide polymorphisms, SNPs) along the 19 chromosomes. Excessively heterozygous sites (false discovery rate, FDR < 0.01) were filtered using BCFtools (Danecek et al. 2021) and low frequency sites (--geno 0.05), and sites with Hardy-Weinberg equilibrium exact test *P*-values below 1e^-6^ (-- hwe 0.000001), were filtered using PLINK (v1.90b4.9, Chang et al. 2015). A total of 6,806,717 bi-allelic SNPs were filtered to minor allele frequency > 5% and annotated by intersection with browser extensible data (BED) file from the genome annotation of the *Populus tremula* reference genome (Schiffthaler et al., 2019) (available at plantgenie.org; Sundell et al., 2015). SNPs were considered as intergenic if they laid further than 2 kbp away from a gene, while SNPs within 2 kbp of a gene were considered associated with that gene. Best DIAMOND matches were used to identify genes in *Arabidopsis thaliana* homologous to the annotated *P. tremula* genes.

Genome wide association mapping was conducted using GEMMA (Zhou and Stephens, 2012) with univariate Linear Mixed Models (LMMs), which are association tests between SNP markers and phenotypic BLUP values. Two covariates were included in the GWAS model: although relatedness in the SwAsp collection was weak, the first covariate was a relatedness matrix of all individuals in the study that was centre-scaled in GEMMA (using the parameter “-gk 1”) as previously described (Wang et al., 2018a); the second was the latitude of origin of each SwAsp genotype, which was applied to eliminate any spurious associations resulting from size differences of the trees that result from the latitudinal sampling cline that influences seasonality-determined growth in the SwAsp collection (Luquez et al., 2008). False discovery rate (FDR) of each association was calculated as the “*q*-value” using R (Storey et al., 2021) following the principle of the Benjamini-Hochberg procedure (Storey and Tibshirani, 2003). The percentage of phenotypic variance explained (PVE) by each SNP, for each trait, was also estimated using the formula described previously (Wang et al., 2018a). GWAS results were visualised using Manhattan plots generated in the ‘qqman’ package in R (Turner, 2017). The distributions of the phenotypic data were tested for normality using Shapiro-Wilk tests, transformed with ordered quartile normalization (described above), and homogeneity of variances tested with a Bartlett test in R prior to analyses of variance amongst SNP genotype groups. Allele boxplots were generated using the ‘ggplots2’ package in R (Wickham, 2016). The ‘anova’ function was applied in R to a linear model where the phenotype was the dependent variable and the SNP allele class the independent variable. Linkage disequilibrium R-squared values were obtained using ‘--r2 with-freqs’ for a list of SNPs defined by ‘--ld-snp-list’ in PLINK (v1.90b4.9, Chang et al. 2015).

## Declarations

### Ethics approval and consent to participate

Not applicable.

### Consent for publication

All authors consent to the publication of this manuscript.

### Availability of data and materials

All the data used for analyses in this manuscript is either displayed in the supplementary datasets, or available upon request to the corresponding author.

### Authors’ contributions

HT designed the study, with help from KMR, SJ, GS, LGS, LJJ, NRS, and SE. Phenotypic characterization of the trees was performed by SE, ML, MLG, KMR, and TG, with supervision by LJJ, GS and HT. OPLS modelling of traits was performed by SE. LGS performed analyses of broad-sense heritability and genetic correlations. Genome-wide association study for identification of SNPs was performed by KMR, NM and NRS. SE and HT wrote the manuscript, with assistance from all co-authors.

## Acknowledgements

The authors thank the UPSC Biopolymer Analytical Platform (supported by Bio4Energy and TC4F) and its manager, Junko Takahashi-Schmidt, for the analyses of the wood chemical composition traits. We thank Veronica Bourquin and Marlene Karlsson for help in preparing the wood samples for analyses, and Daria Chrobok for the illustration in Fig. 2. We thank Skogforsk at Ekebo for hosting the SwAsp common garden and Magnus Alsterfjord for help with the field sampling.

## Supplementary information

**Additional file 1**. The median value of each trait for each of the SwAsp *Populus* lines and correlation of each trait with the latitude of genotype origin.

**Additional file 2**. Statistical estimation of broad-sense heritability H^2^ and genetic coefficient of variation CV(G) for all the monitored traits.

**Additional file 3**. Genetic correlations between the tree growth, wood and saccharification traits.

**Additional file 4**. Significant (*q*-value <0.1) single nucleotide polymorphisms (SNPs) identified in the genome-wide association study (GWAS) of 65 traits recorded in the Swedish aspen common garden.

