## Additional file 2 for "Genetic markers and tree properties predicting wood biorefining potential in aspen (*Populus tremula*) bioenergy feedstock"

**Additional file 2. Statistical estimation of broad-sense heritability ($H^2$) and genetic coefficient of variation CV(G) for all the monitored traits.** Full description of the traits can be found in the Additional file 1.

| Trait full name | Trait abbreviation | $H^2$ | standard error for $H^2$ | CV(G), % |
|---|---|---|---|---|
| Arabinose unit content | Ara | 0.08 | 0.04 | 7 |
| Arabinose release in hydrolysate after pretreatment | ARA_EH_PT | 0.00 | 0.00 | 0 |
| Arabinose release in pretreatment liquid | ARA_PL_PT | 0.21 | 0.06 | 8 |
| Arabinose release without pretreatment | ARA_UT | 0.05 | 0.04 | 7 |
| Carbohydrate content | C | 0.27 | 0.05 | 1 |
| Carbohydrate to lignin ratio | CL | 0.26 | 0.05 | 5 |
| Coarseness of the wood | Coarseness | 0.74 | 0.03 | 9 |
| Wood density | Dens | 0.59 | 0.04 | 8 |
| Wood density, excluding vessels | DensVessFre | 0.66 | 0.04 | 9 |
| Stem diameter at breast height | DBH | 0.75 | 0.03 | 39 |
| Tree height | Height | 0.79 | 0.03 | 29 |
| Fraction of the wood made of fibers | FibFrac | 0.66 | 0.04 | 3 |
| Fiber cell perimeter | FibPerim | 0.67 | 0.04 | 4 |
| Ratio of fiber to vessel cells | FibPerVess | 0.55 | 0.05 | 17 |
| Fiber population | FibPop | 0.64 | 0.04 | 8 |
| Fiber radial diameter | FibRadDia | 0.69 | 0.04 | 5 |
| Fiber tangential diameter | FibTanDia | 0.64 | 0.04 | 3 |
| Fucose unit content | Fuc | 0.43 | 0.05 | 13 |
| G-lignin content | G | 0.54 | 0.05 | 8 |
| Galactose unit content | Gal | 0.08 | 0.05 | 6 |
| Galacturonic acid unit content | GalA | 0.08 | 0.04 | 6 |
| Galactose release in hydrolysate after pretreatment | GAL_EH_PT | 0.00 | 0.00 | 0 |
| Galactose release in pretreatment liquid | GAL_PL_PT | 0.11 | 0.05 | 7 |
| Galactose release without pretreatment | GAL_UT | 0.21 | 0.05 | 8 |
| Glucose unit (from non-crystalline cellulose) content | Glc | 0.14 | 0.05 | 8 |
| Glucuronic acid unit content | GlcA | 0.56 | 0.05 | 41 |
| Glucose release in hydrolysate after pretreatment | GLU_EH_PT | 0.17 | 0.05 | 3 |
| Glucose release in pretreatment liquid | GLU_PL_PT | 0.11 | 0.05 | 6 |
| Glucose release without pretreatment | GLU_UT | 0.26 | 0.05 | 17 |
| Glucose production rate after pretreatment | GPR_PT | 0.22 | 0.05 | 6 |
| Glucose production rate without pretreatment | GPR_UT | 0.20 | 0.05 | 11 |
| H-lignin content | H | 0.40 | 0.06 | 22 |
| Total hexose release after pretreatment | HEX_PT | 0.12 | 0.05 | 3 |
| Total lignin content | L | 0.29 | 0.05 | 4 |
| Mannose unit content | Man | 0.39 | 0.05 | 19 |
| Mannose release in hydrolysate after pretreatment | MAN_EH_PT | 0.17 | 0.05 | 9 |
| Mannose release in pretreatment liquid | MAN_PL_PT | 0.24 | 0.05 | 11 |
| Mannose release without pretreatment | MAN_UT | 0.28 | 0.05 | 12 |
| Microfibril angle | MFA | 0.49 | 0.05 | 16 |
| Modulus of elasticity | MOE | 0.52 | 0.05 | 14 |
| 4-O-methyl-glucuronic acid unit content | 4-O-meGlcA | 0.27 | 0.05 | 8 |

| | | | | |
|---|---|---|---|---|
| Phenolics content excluding S-, G- and H-lignin | P | 0.34 | 0.05 | 6 |
| Total pentose release after pretreatment | PENT_PT | 0.01 | 0.04 | 2 |
| Rhamnose unit content | Rha | 0.19 | 0.05 | 6 |
| S-lignin content | S | 0.43 | 0.05 | 6 |
| Ratio of S-lignin to G-lignin content | SG | 0.72 | 0.04 | 11 |
| Total arabinose released after pretreatment (in pretreatment liquid and in hydrolysate) | Total_ARA_PL+EH_PT | 0.21 | 0.05 | 8 |
| Total galactose released after pretreatment (in pretreatment liquid and in hydrolysate) | Total_GAL_PL+EH_PT | 0.11 | 0.05 | 6 |
| Total glucose released after pretreatment (in pretreatment liquid and in hydrolysate) | Total_GLU_PL+EH_PT | 0.15 | 0.05 | 3 |
| Total mannose released after pretreatment (in pretreatment liquid and in hydrolysate) | Total_MAN_PL+EH_PT | 0.25 | 0.05 | 10 |
| Total sugar released after pretreatment | TOTAL_SUGAR_PT | 0.02 | 0.04 | 1 |
| Total xylose released after pretreatment (in pretreatment liquid and in hydrolysate) | Total_XYL_PL+EH_PT | 0.01 | 0.04 | 2 |
| Unkown cell wall compounds content | U | 0.15 | 0.05 | 3 |
| Cell wall thickness of wood cells | Wall_thickness | 0.69 | 0.04 | 9 |
| Fraction of the wood made of vessels | VessFrac | 0.66 | 0.04 | 11 |
| Vessel major diameter | VessMajorDiam | 0.64 | 0.04 | 10 |
| Vessel minor diameter | VessMinorDiam | 0.63 | 0.04 | 8 |
| Vessel perimeter | VessPerim | 0.65 | 0.04 | 9 |
| Vessel population | VessPop | 0.52 | 0.05 | 16 |
| Unidentified cell wall compounds content | 0 | 0.12 | 0.05 | 2 |
| Xylose unit content | Xyl | 0.05 | 0.04 | 2 |
| Xylose release in hydrolysate after pretreatment | XYL_EH_PT | 0.12 | 0.05 | 8 |
| Xylose release in pretreatment liquid | XYL_PL_PT | 0.01 | 0.04 | 2 |
| Xylose release without pretreatment | XYL_UT | 0.11 | 0.05 | 16 |
